# Sensations from a single M-cone depend on the activity of surrounding S-cones

**DOI:** 10.1101/260653

**Authors:** Brian P. Schmidt, Ramkumar Sabesan, William S. Tuten, Jay Neitz, Austin Roorda

**Affiliations:** Graduate Program in Neuroscience, University of Washington, Seattle WA, 98109, USA; School of Optometry and Vision Science Graduate Group, University of California, Berkeley CA, 94720, USA; Department of Ophthalmology, University of Washington, Seattle WA, 98109, USA

**Author notes:** These authors contributed equally (Brian P. Schmidt, Ramkumar Sabesan). brian (Brian P. Schmidt).

## Abstract

Color vision requires the activity of cone photoreceptors to be compared in post-receptoral circuitry. Decades of psychophysical measurements have quantified the nature of these comparative interactions on a coarse scale. How such findings generalize to a cellular scale remains unclear. To answer that question, we quantified the influence of surrounding light on the appearance of spots targeted to individual cones. The eye’s aberrations were corrected with adaptive optics and retinal position was precisely tracked in real-time to compensate for natural movement. Subjects reported the color appearance of each spot. A majority of L-and M-cones consistently gave rise to the sensation of white, while a smaller group repeatedly elicited hue sensations. When blue sensations were reported they were more likely mediated by M- than L-cones. Blue sensations were elicited from M-cones against a short-wavelength light that preferentially elevated the quantal catch in surrounding S-cones, while stimulation of the same cones against a white background elicited green sensations. In one of two subjects, proximity to S-cones increased the probability of blue reports when M-cones were probed. We propose that M-cone increments excited both green and blue opponent pathways, but the relative activity of neighboring cones favored one pathway over the other.

## Introduction

Human trichromatic color vision is mediated by three classes of cones: long (L) medium (M) and short (S) wavelength sensitive^1,2^. Individually, cone photoreceptors are colorblind^3^. To extract color information from the cone mosaic, the visual system compares the relative rate of photon absorptions across the three cone types^4^. Over many decades, color scientists have measured the contribution of each cone type to the chromatically opponent red-green and blue-yellow pathways using carefully tailored large field stimuli^5^. The linking hypothesis underlying this effort was that there exist spectrally opponent neurons with cone interactions that match those measured psychophysically and thus provide a physiological substrate for low-level color appearance judgments^6^. However, the effort to unify the physiology of the primate visual system with color perception has proven difficult. The major spectrally-opponent channels found in the lateral geniculate nucleus, for instance, do not map directly onto hue opponent channels measured psychophysically^7^.

A related technique for inferring the properties of these post-receptoral interactions is the recording of percepts elicited from small, dim flashes of light^8–14^. The number of cones activated by a near-threshold small spot can be restricted to as few as one or two. This paradigm has contributed, for example, to the discovery of foveala tritanopia^15,16^, the spatial arrangement of L- and M-cones across the retina^17–19^ and the role of M-cones in the perception of blue^20^. The appearance of small spots also raises questions about how well the large field studies generalize to cellular scales. For example, the color sensations elicited from cone-sized spots differ even between spots absorbed by cones with the same photopigment^10^. Presumably, understanding these within cone-class variations in color appearance will help to reconcile perception with single-cell physiology. However, prior studies using this tool have been restricted by an inability to spatially target light to identified photoreceptors.

Recently, we used adaptive optics with high-resolution eye-tracking technology^21,22^ to stimulate cones of known spectral type^23^ and revealed the first cellular-scale topographic map of color appearance. Our results demonstrated that in the presence of an unchanging, white background, each L- or M-cone predominantly elicited a single color percept. However, unlike the results of Hofer et. al.^10^ in which subjects reported blue percepts even though their lights did not stimulate S-cones, under our conditions subjects used only three color terms – red, green and white – to describe their experience of cone-targeted flashes. We hypothesized that the specific color terms used were a consequence of the baseline activity in the cones surrounding those being targeted, since the percept elicited from stimulation of a single cone will depend upon the activity in neighboring cones as well. It follows then that the sensations would change if the same cone was targeted against a different context of baseline activity in surrounding cones^12^ and the nature of this change will provide insight into how color information is extracted from the cone mosaic.

In the present work, a short-wavelength dominant background that preferentially increased S-cone activity was used and color reports of small spots targeted to single cones were collected. Under these conditions, subjects frequently reported blue percepts when M-cones were targeted. These findings imply that the relative activity in S-cones can, in some cases, influence the appearance of light targeted to other cones. We speculate about the neural pathways that may mediate this behavior in the Discussion.

## Results

### The effect of elevating the activity of surrounding S-cones on color naming

Cones were targeted with small spots of light (543 nm) against a short-wavelength dominant background in two male subjects. Sensitivity to 543 nm is similar for L- and M-cones. After each stimulus, color naming responses were collected. Previously, we reported color naming results with the same apparatus, subjects and stimulus conditions, but against a white background^23^.Figure 1A demonstrates the approximate appearance of large field L- and M-cone isolating stimuli against a blue background. In this context, an M-cone selective increment yields a blue hue. In contrast, an M-cone increment relative to a white background appears green. The dominant hue in the case of L-cone stimuli is red in both cases. It is not known whether changing background contexts will have the same phenomenological influence on single cone stimuli. Targeting light to a single cone produces far higher cone-contrast than traditional experimental methods. Additionally, the restricted nature of a stimulus that excites a single cone may activate only a subset of pathways invoked during natural viewing.

**Figure 1.**
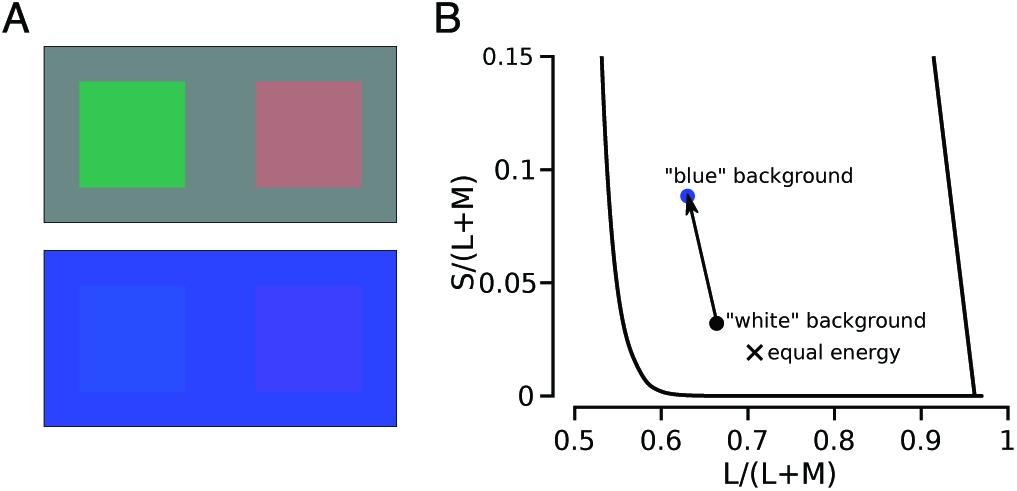
Experimental design. (**A**) Simulations of the white (upper panel) and blue backgrounds (lower panel) adopted in our experiments are shown. The four squares represent large-field cone-isolating stimuli (~9% cone-contrast). The right squares are stimuli that selectively increase the quantal catch only in L-cones compared to the rectangular backgrounds. The left square stimuli selectively increase the quantal catch in M-cones. Relative to a gray background, the M-cone isolating stimulus appears greenish or greenish-blue, while the L-isolating stimulus looks red or pink. Relative to a blue background, the stimuli appear blue and reddish-blue, respectively. (**B**) The chromaticity of each background condition is shown^56^. Black circle = white, blue circle = blue background used in the experiment; black ‘X’ denotes ‘equal energy’. In this chromaticity space, change in the y-dimension (S/(L+M)) indicates an isolated change in S-cone activity. Relative to the white condition, S-cone activity (y-axis) was increased in the blue context with a smaller change in M:L-cone activity (x-axis).

The blue background *increased* the quantal catch of S-cones by a factor of 2.35 relative to the white background. L-and M-cones were altered less so: activity *decreased* by factors of 1.24 and 1.07 in L- and M-cones, respectively (Figure 1). Changing the background to blue did not alter the subject’s ability to see the stimulus and frequency of seeing was similar between L- and M-cones (Figure 1C) (both backgrounds yielded 88% seen). The observation that L- and M-cone frequency of seeing was comparable in both background conditions indicates that adaptation within these cone sub-types did not substantially influence their sensitivity. While we did not carry out formal threshold measurements and subjects were free to repeat trials, we postulate that the intensity of each flash, which was much greater than the background intensity, minimized the influence of the background field on sensitivity in L- and M-cones.

A total of 258 L/M-cones were targeted under the blue condition. S-cones were insensitive to the stimulation wavelength. To permit a detailed comparison with the previously reported white background data, we targeted many of the same cones on the blue background (N=246). To assess the reliability of percepts, the same cone (N=51) was often targeted during more than one session. When the responses obtained during separate sessions – which often occurred weeks or months later – were compared, we found that the responses associated with a given cone were repeatable; agreement=72.4 ±3.0%. The percepts recorded with this technique were also previously shown to be largely invariant to changes in wavelength and fluctuations in intensity^23^.

The presence of a blue background altered color naming relative to the previously tested white condition. Against a white context, a majority of trials produced achromatic reports, while a minority yielded red or green percepts. Here, against a blue background, a majority of flashes were again labeled achromatic and a smaller fraction red. However, both subjects ceased to call flashes green and instead used blue to describe a fraction of trials. Both subjects verbally reported that some blue sensations were tinged with green, but the magnitude and frequency of this phenomenon was not quantified. Similar to a white context, our two subjects never used yellow to describe the appearance of flashes, which may be related to the relative difficulty of discriminating a white from a yellow spot^14^. Below we consider the influence of a blue background on single cone color reports in more detail.

### M-cones contribute to blue pathways

The single largest predictor of color percept was the spectral class (L- or M-) targeted. Previously, against a white context, we observed that L- and M-cone trials were equally likely to give rise to a white sensation, but L-cones more frequently mediated red reports and M-cones were more likely to produce green sensations^23^. Against a blue background, the average L- (65.8±1.9%) and M-cone (61.4±2.6) again produced a similar number of white reports (Figure 2A). However, on any given trial a white report was slightly more likely when an L-cone was targeted: Fisher’s Exact odds ratio=1.18, *p* < 0.001. Red percepts were more likely to be driven by L- (21.3±2.0%) than M-cones (4.2±1.1%): Fisher’s Exact odds ratio=5.88, *p* < 0.001. Blue percepts, on the other hand, were more likely to be mediated by M- (34.4±2.9%) than L-cones (12.9±1.7%): Fisher’s Exact odds ratio = 3.34, *p* < 0.001. The latter observation is consistent with the interpretation that M-cones mediated blue sensations in previous small spot experiments^9,10,12,20,24^.

**Figure 2.**
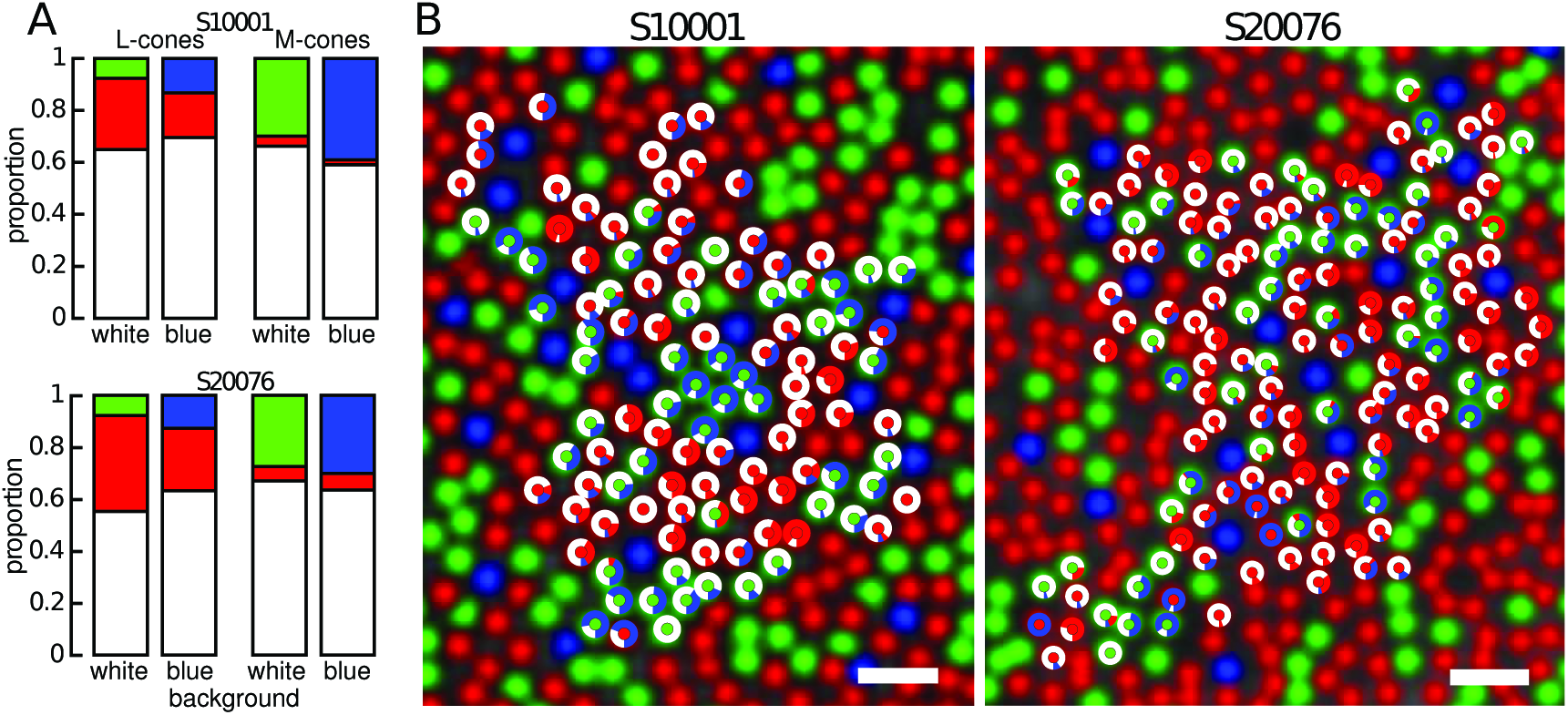
Color appearance of cone targeted stimuli on a blue background. Color naming distributions from S10001 and S20076. (**A**) Proportion of responses after stimulation of L- and M-cones on a white and blue background. (**B**) The response to stimulation with a 543 nm light against a blue background is represented for each cone by a donut plot. The center of the donut indicates the type of cone targeted (L=red, M=green). Colors correspond to reported percepts. Scale bar = 2.5 arcmin.

Next, we investigated the relationship between spectral type and background on a cone-by-cone basis. Correlation analyses revealed that color reports on one background were highly predictive of reports on the other (in all comparisons *R* > 0.48, *p* < 0.001). This trend is illustrated by Figure 3, which displays the color naming data transformed into a two-dimensional representation of red-green and blue-yellow. Cones that, on average, produced a white sensation lie at the origin of this plot. Each cone is plotted twice to represent its behavior on both backgrounds and is joined by a solid line to show how the background influenced the cone’s behavior. The diagonal lines connecting M-cones confirm that they switched from mediating green to blue sensations and the cones that strongly signaled hue tended to do so in both contexts. A small number of L-cones also exhibited this behavior. Most L-cones fell along the white-red axis, with only occasional blue responses. Together, these results demonstrate the substantial influence the relative activity of S-cones has on the color appearance of small spots of light.

**Figure 3.**
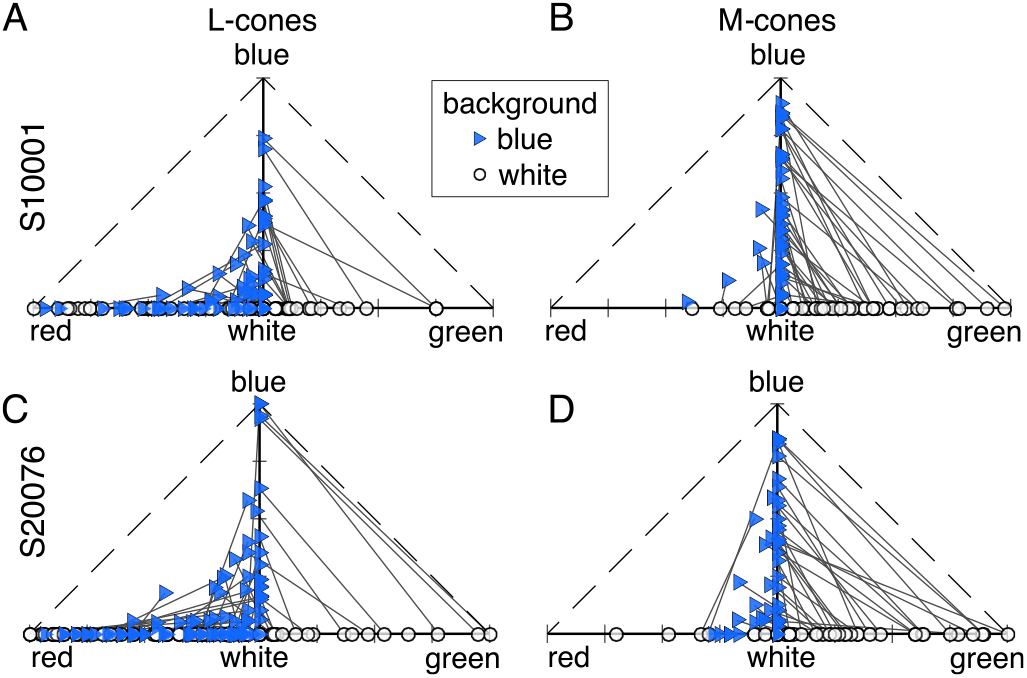
Opponent color responses from a population of L- and M-cones. Color distributions for each cone were transformed into a red-green 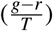 and a blue-yellow 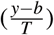 dimension, where *r, g, b, y* are the number of trials that produced red, green, blue and yellow responses, respectively and *T* equals the total number of trials. Each marker represents a cone that was targeted on both a white (circles) and blue (triangles) background. Left column = L-cones, right column = M-cones. Each cone is plotted twice and connected by a solid line to represent its color distribution in both contexts. **A&B** display the data from S10001 and **C&D** represent S20076. Yellow dimension is not shown because subjects never used yellow in these experiments.

### Nearby cones predict color responses

The local neighborhood surrounding a cone has been previously hypothesized to play a role in determining color percepts^19,25–28^. To test this hypothesis, we determined the six nearest neighbors to each cone and analyzed the influence of this local neighborhood on color reports. In Figure 4 we plotted the proportion of red, green and blue responses as a function of nearest neighbors that were of a different spectral class. In 4 out of 6 cases the effects of the local neighborhood were statistically significant (Figure 4A, B, D and F). However, the strength of correlation was weak in all cases (*R*^2^ = 0.18). The most substantial effect of neighborhood was on the proportion of blue responses elicited upon stimulation of M-cones. We found a weak, but significant tendency for M-cones to signal blue when they were surrounded by other M-cones (N=99, *R*^2^ = 0.18, *p* < 0.001), which contradicts the predictions of some models of color opponency^19,25–28^. In all cases, the influence of nearby neighbors decreased when the analysis was extended to cones beyond the nearest six, because including cones beyond the nearest six yielded neighborhoods that were increasingly similar to the L:M:S ratio over the whole mosaic and, therefore, captured less of the variance in color naming. Analogous results were obtained when we ran the same analyses on color naming reports from the white background condition^23^.

**Figure 4.**
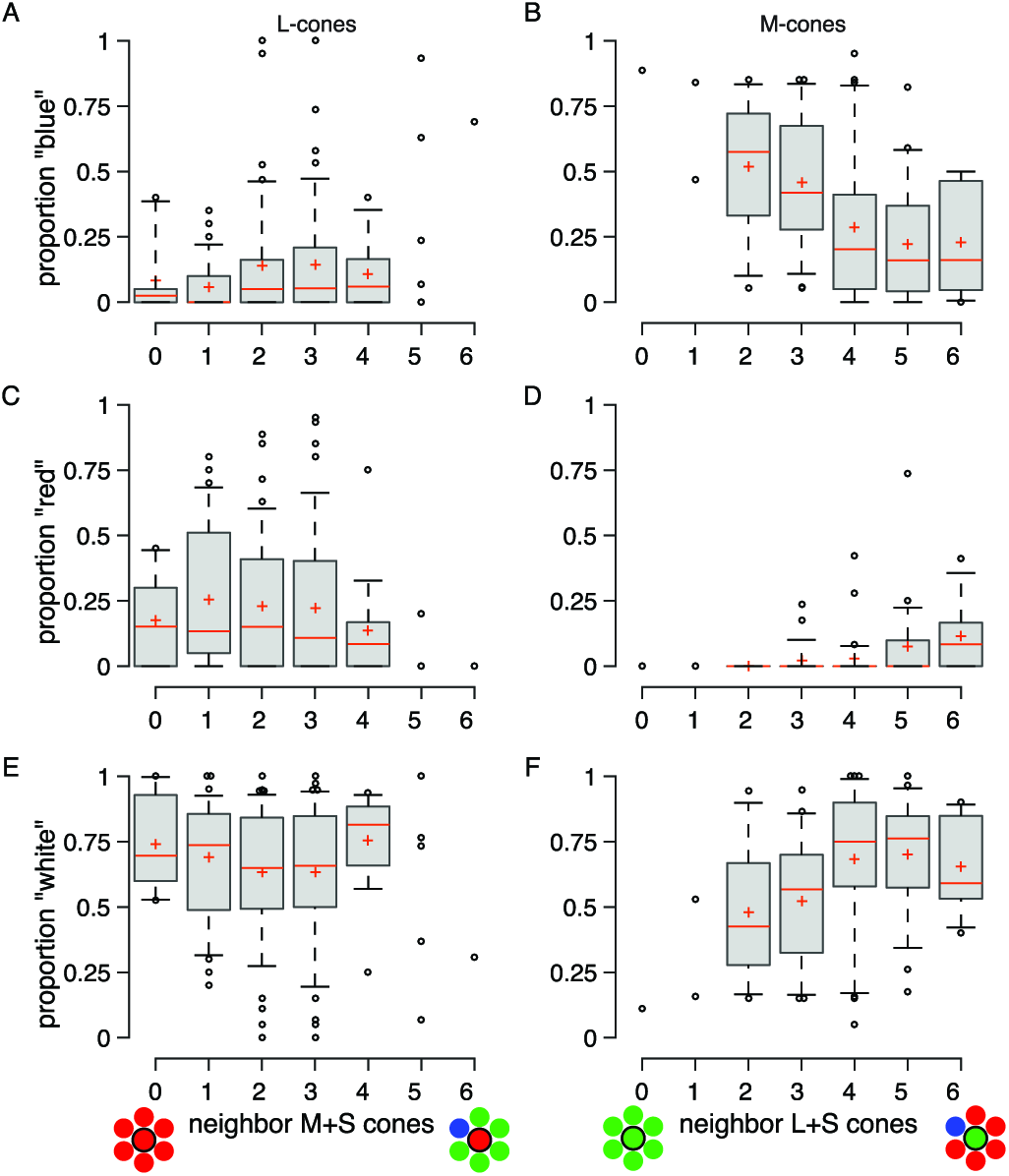
Influence of the local cone mosaic on color percepts. Box-and-whisker plots represent the proportion of responses named blue, red and white for L- (left column; N=159) or M-cones (right column; N=99) stimulated against a blue background. The proportion of blue (**A&B**), red (**C&D**) and white (**E&F**) is represented as a function of each cones’ neighborhood. Red lines represent the median, crosses the mean; boxes span the 25th and 75th percentiles; whiskers span 9 and 91%. Spearman rank correlation coefficients: **A**. *R* = 0.199, *p* = 0.012; **B**. *R* = –0.419, *p* = 2 × 10^−5^; **C**. *R* = –0.14, *p* = 0.078; **D**. *R* = 0.371, *p* = 0.0002; **E**. *R* = –0.025, *p* = 0.752; **F**. *R* = 0.325, *p* = 0.001.

Another potential influence on color percepts is the presence of S-cones in the local neighborhood. In one subject, we found evidence for a dependence on the distance to nearby S-cones. Figure 5 displays the relationship between the mean distance to the nearest three S-cones and the proportion of responses named blue when M-cones were targeted. In S10001, we found a greater proportion of blue reports when an M-cone was in close proximity to S-cones (N=48, *R* = –0.487, *p* < 0.001). The distance to the nearest three S-cones was chosen because S-cones are relatively scarce in the mosaic and their signals presumably sum in post-receptoral circuits. Regardless, the distance to the single or two nearest S-cones also produced statistically significant (*p* < 0.05) predictions of blue responses in this subject. However, we did not observe the same trend in our second subject (N=51, *R* = –0.008, *p* = 0.96). Because this analysis is sensitive to the presence or absence of S-cones, one possible reason for this discrepancy maybe the mis-classification of just a few S-cones by our densitometry method^29^.

**Figure 5.**
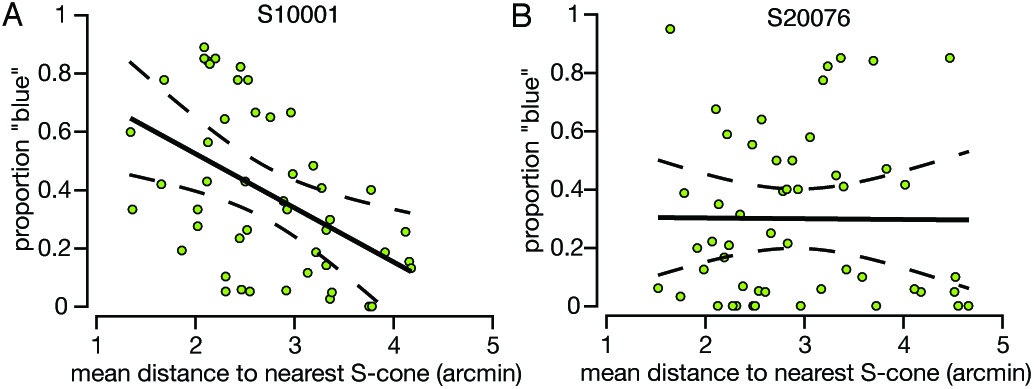
Dependence of blue percepts on nearby S-cones. The proportion of blue responses when an M-cone was stimulated is plotted as a function of mean distance to the three nearest S-cones. The proportion of blue responses from L-cones was not significantly correlated with S-cones. Each point in the plot represents an M-cone that was targeted on a blue background. The solid line represents the best-fit linear regression; dashed lines indicate 95% confidence intervals. (**a**) Subject S10001. Regression equation: *y* = –0.186*x* + 0.896. (**b**) Subject 20076. Regression equation: *y* = –0.003*x* + 0.308.

Finally, the predictors themselves may be correlated since both the distance to the nearest S-cone (Figure 5) and the number of non-like neighbors (Figure 4) are dependent upon the spatial layout of the mosaic. To test this possibility, we ran a multiple regression that included both predictors of blue sensations from M-cones. The analysis was significant for data collected from both subjects (S10001: N=48, *R* = 0.613, F_45_=13.57, *p* = 2 × 10^−5^; S20067: N=51, *R* = 0.354, F_48_=3.44, *p* = 0.04). Post-hoc analyses for S10001 revealed that both neighborhood (*t* = –3.17, *p* = 0.003) and S-cone distance (*t* = –3.32, *p* = 0.002) contributed significantly to the regression. In S20076, the only significant predictor was the number of opponent neighbors (*t* = –2.62, *p* = 0.01).

### Discussion

In these studies, examining the perception of cone-targeted spots, we found that preferentially elevating the activity of surrounding S-cones enhanced the perception of blue in flashes absorbed by individual M-cones. The changes in hue reports demonstrate that light absorbed by nearby cones can influence the percept assigned to the most elementary units of visual input.

The appearance of large spots, which stimulate many cones, are often influenced by the chromaticity of the surrounding field. This phenomenon, known as color induction^30^ includes both assimilation, when the color of a spot shifts towards the background, and chromatic contrast, when the appearance of a spot shifts away. In the presence of a blue background like the one used here, chromatic contrast would induce yellowness into the test spot. However, subjects never saw yellow. Rather, subjects reported blue sensations on 21% of trials, which was was consistent with color assimilation. Assimilation has been explained at a neural level by assuming that chromatic neurons with both small and large receptive fields sample the same area of visual space in parallel^30,31^. Color-opponent neurons with large receptive fields will carry a spatially-averaged chromatic signal (*i.e*. the mean of the spot plus background). Meanwhile, cells with small receptive fields will be more acutely sensitive to the chromaticity of the small spot. If these parallel streams are combined in cortical neurons, which sum inputs from both types of cells, then their mixed signal may retain the resolution of the small cells but contain a color bias towards the background. We cannot exclude the possibility that assimilation influenced, to some extent, the appearance of stimuli in our study. For instance, many L-cones that yielded red or white reports on a majority of trials against a white background, occasionally elicited blue reports when the background was blue (Figure 2B). However, assimilation cannot entirely explain why only a small group of M-cones produced a majority of blue sensations or how a majority of L/M-cones yielded mostly white reports against both a white and a blue background. Rather, these results suggest that the background activity of neighboring cones influenced the response of color cells with relatively small receptive fields, as well as those with larger receptive fields. Below we consider possible mechanisms for this influence.

The blue background in our study produced a large change in S-cone quantal catch (an increase of 135% over the white condition, *i.e*. 2.35 times) and a much smaller change in the ratio of M:L activity (16% increase, *i.e*. 1.16 times) relative to the previously tested white background (Figure 1). The experimental conditions were otherwise identical to our prior work^23^. Against the white context, M-cones that mediated a hue overwhelmingly produced green sensations. After elevating activity of S-cones, M-cones gave rise to blue, rather than green, percepts (Figure 2) and the cones that gave rise to hue sensations were the same ones most likely to mediate a hue sensation against the white context (Figure 3). On the other hand, additional factors, like the spectral neighborhood of a targeted cone, had comparatively little influence on the color perceived (Figure 4). It has been previously observed that small, middle-wavelength spots are sometimes perceived as blue. Because S-cones are relatively insensitive to wavelengths beyond 500 nm, it has been proposed that such “blue” flashes of light were detected by M-cones^9,10,12,20,27,32^. The current study directly confirmed that proposal; when increments of light were spatially confined to identified M-cones the spots often appeared blue. In our case, this happened when we elevated the quantal catch of surrounding S-cones.

Having confirmed that M cones mediate blue sensations, the question becomes how is it that a wavelength of light (543 nm) that does not stimulate S-cones elicits blue sensations when targeted to single M-cones? Hofer et al.^10^ suggested that activity in a single M-cone mimics the ratio of excitation among the three cone classes that occurs when we normally perceive blue. For instance, a 475 nm monochromatic light, which appears to the average observer as purely blue, produces quantal catches in the S:M:L cones with a ratio of 47:31:18^33^. The corresponding ratio for a pure green 520 nm light is 1.8:89:68. Hofer et al.^10^, noted that, compared to green lights, blue lights stimulate M cones more strongly than L. They suggested that perhaps M-cones gave rise to blue sensation under their conditions because targeting a single M-cone excited it much more strongly than surrounding L-cones, thus mimicking the high M:L excitation ratio that is more associated with blue light. In the present work, when an M-cone was targeted the ratio of M-cone activity elicited by the spot relative to L-cone excitation in the surround was high and approximately the same between conditions. Thus, M:L-cone activity alone was not likely to be the factor that determined when M-cones signaled blue. A second major difference between conditions that normally produce blue vs. green sensations is that the S-cone quantal catch, according to the ratios above, is about 26 times higher (47/1.8) for pure blue than pure green and our blue background condition may mimic that by increasing the quantal catch of surrounding S-cones. According to color opponent theory, blue is seen when there is activity in a blue-yellow channel and green appears when there is activity in a green-red channel. When both channels are active, a mixture of blue and green is seen, which reflects the relative activity in these two channels. M-cone increments are thought excite the sensation of green via an M – (S+L) pathway and blue via an (S+M) – L opponent pathway. We propose that the elevated activity in S-cones biased post-receptoral pathways with summed input from S- and M-cones. By elevating the activity of S-cones, we suppressed the green pathway and excited the blue mechanism.

We now turn to the question of which post-receptoral pathway most likely mediated this shift. As noted above, a candidate pathway must be (1) color opponent and (2) have small receptive fields that will be excited by a cone-sized stimulus. In the primate retina, the three smallest receptive fields, in order of increasing size, belong to the parvocellular (midget ganglion cells), magnocellular (parasol ganglion cells) and koniocellular (small bistratified ganglion cell) pathways^34^. Near the fovea where our experiments were conducted, the center mechanism of parvocellular neurons draws input from a single cone^35^ and carries L-M cone opponent signals^34^. For these reasons, they have been identified as candidates for carrying information about both color vision and high-resolution spatial details^36–40^. Parvocellular neurons are also known to respond to our stimuli. Using a similar system to the one used here, Sincich et al.^41^ found robust and repeatable responses in peri-foveal (3.7-6°) parvocellular neurons stimulated with cone-sized spots. Neurons in the magnocellular pathway may also be excited by a cone-targeted stimulus^42,43^. However, magnocellular neurons do not carry a chromatic signal^44^. Evidence from both behavioral^45^ and recent *ex vivo* physiological investigations^46^ suggests that these neurons carry information most critical for motion processing. Finally, the koniocellular pathway carries an S-ON/LM-OFF spectrally opponent signal and is more commonly associated with blue-yellow color vision^47,48^. However, the incremental stimuli in our study are not ideal for eliciting activity from this pathway. L/M-cones provide inhibitory input to this channel and thus targeting them individually ought to decrease the sensation of blue and increase yellowness – a prediction that does not square with our findings. On the other hand, the short-wavelength background in our study very likely increased activity in the S-ON/LM-OFF pathway, which may have contributed to the large field component of assimilation.

Thus, the parvocellular pathway was most likely to have mediated the observed small field color changes. If S-cones contribute excitatory input to an OFF M-L pathway and inhibitory input to an ON M-L parvocellular pathway, they would form an (S+M)-L circuit for blueness and an M-(L+S) circuit for greenness^36,37,49^. In this model, hyperpolarization of S-cones would favor one pathway over the other and explain the observed shifts in hue perception reported here. One potential location where S-cone signals could be summed with L/M opponent pathways is in the outer retina^50^, where cones laterally influence the output of nearby receptors. Support for this site of lateral interaction was observed in the relationship between proximity to S-cones and the M-cones most likely to mediate a blue percept (Figure 5). Recent threshold measurements of cone-targeted stimuli have also revealed small-scale lateral interactions between S- and L/M-cones, which were also consistent with a retinal origin^51^. Alternatively, S-cone signals could be introduced to M-L opponent parvocellular neurons in cortical circuits^27^. Cortical neurons carrying both (S+M)-L opponency have been reported, but it is not known where in the visual pathway the S-cone input is first introduced^52^.

The methods described here make it possible to direct small spots of light to individual cones of known spectral type. By recording the sensations perceived from stimulation of individual cones, we related the spectral arrangement of the cone mosaic to the private experience of color. The results described here provide evidence for fine scale spatial interaction between S-cones and M-cones that shapes the color appearance of small flashes of light.

## Methods

### Subjects

Two highly experienced adult male Caucasian subjects (S10001 and S20076) participated in the study. Both were authors of the study, had normal color vision and participated in a related study on a white background^23^. Prior to experiments, drops of 1% tropicamide and 2.5% phenylephrine were administered to dilate the pupil and paralyze accommodation. Bite bars were used to stabilize the subject. Informed consent was obtained from each subject and all experimental procedures adhered to the tenets of the Declaration of Helsinki and were approved by the University of California Berkeley Institutional Review Board.

### Cone-resolved imaging and stimulation

An adaptive optics scanning laser ophthalmoscope (AOSLO) was used to image and stimulate individual cones. The details of the experimental apparatus and the procedure for targeted micro-stimulation have been previously reported in detail^21–23^. Briefly, the paradigm utilized an 840 nm (24 nm full-width at half maximum bandwidth) infrared (IR) channel for imaging and a 543 nm (22 nm full-width at half maximum bandwidth) channel for stimulation. The 543 nm wavelength was chosen to minimize the difference in L- and M-cone activity. Both wavelengths were raster scanned to generate a 1° imaging field of view on the retina. The 840 nm light was used to both measure optical aberrations (10%) and image the retina (90%). Optical aberrations were measured with a wavefront sensor. The data from the wavefront sensor was subsequently used in real-time to control a deformable mirror that corrected for the measured aberrations. The resulting correction produced diffraction-limited imaging and stimulus delivery. The remaining 840 nm light was collected in a photo-multiplier tube through a confocal pinhole and rendered into a video stream that was used to perform real-time high-resolution image registration. Retinal tracking coordinates obtained from this procedure were used drive an acousto-optical modulator; a high-speed (50 MHz) optical switch delivering 543 nm visual stimuli to the retina one pixel at a time. The precision of the procedure was previously measured to be 0.15 arcmin^22,23^. Longitudinal chromatic aberration between the 543 nm and 840 nm wavelengths was corrected by adjusting the relative vergence of the two channels. Finally, transverse chromatic aberration, the spatial offset between the imaging and stimulating channel, was monitored and corrected following previously published procedures^53^.

### Identifying the spectral type of each cone

Prior to the color naming experiments, the spectral identity (L, M, S) of a mosaic of cones located about 1.5° from the fovea was identified in both subjects. The cone identities in a ~0.5° patch, comprising 800 and 631 cones respectively for S10001 and S20076, were obtained. Detailed densitometry results from S10001 were reported in^29^; S10001 was their subject 1. The percentage of L- cones (L / (L+M)) was 66.2 %L and 66.7 %L, respectively, in S10001 and S20076. The spectral type of each cone was identified using AOSLO imaging described above in conjunction with densitometry. Details of the classification technique have been described elsewhere^29^. Each cone tested psychophysically was identified with a specific spectral type by aligning the 840 nm reflectance image and the corresponding trichromatic cone mosaic map obtained from densitometry.

### Cone-targeted color naming

Each session began by collecting a high signal-to-noise ratio retinal image, averaged from 90 raw AOSLO image frames. The experimenter then marked the cones of interest on that image. Effort was made to select cones contiguous to previously tested areas and to repeat a handful of previously tested cones during each session. The subject initiated each trial with a key stroke. During stimulus presentation, a one-second retinal video was recorded and stimulus onset was denoted by an audible beep. The subject then selected a color name to describe the flash from six possible choices: red, green, blue, yellow, white and not seen. Additional categories were provided during pilot studies, but were not used by the subjects (see below). Each response was registered by a second keystroke, which was accompanied by another audible beep of a different pitch. The time-stamp of optical switch activation was embedded into the retinal video with a digital mark. This mark was retrieved off-line using image cross-correlation and compared against the desired location to arrive at an estimate of spatial offset errors in stimulus presentation for each frame. A subset of cones analyzed with this method yielded average delivery errors of 0.21 (N=94, standard deviation=0.07) and 0.16 (N=149, standard deviation=0.05) arcmin for S10001 and S20076, respectively.

Flashes were 3 pixels (~0.45 arcmin) in diameter and 500 ms in duration. The size of the stimulus was less than half the diameter of a cone inner segment (~1 arcmin) at the retinal eccentricity studied (1.5°). The luminance of the 543 nm stimulus (3570 cd m^−2^) was approximately 80-90 times higher than the background. The power of the stimulus channel was measured with a radiometer (Newport Optical Power Meter 1830-C; Irvine, CA, USA) and converted into quanta. A single (3×3 pixel, 15 frames) flash delivered approximately 7.0 log10 quanta to the cornea.

The color naming technique we adopted required subjects to break up color space into five categories: red, green, blue, yellow and white. Subjects were not given specific instructions on how to judge the color, but instead were encouraged to develop internal category boundaries. Both subjects felt comfortable describing the flashes with these color names. Each subject was given numerous days of practice trials and was only permitted to begin experimental data collection after they felt confident in their category boundaries. Feedback was given after the completion of sessions on the repeatability of cones that were targeted across sessions. Practice continued until repeated sessions of the same cones reached a minimum threshold of *R* > 0.70.

We cannot exclude the possibility that a richer palette of descriptors would have altered the results. For instance, both subjects verbally indicated that some blue percepts appeared tinged with green, which would be consistent with the idea that some cones contribute to both a blue and green pathway that is biased to a greater or lesser extent by modulating S-cone activity. However, the magnitude of bluish-green reports was not quantified. It also remains possible that some white flashes were tinged with a hue, perhaps in some cases yellow, but due to the crudeness of the color categories were lumped in with the white category. For instance, white, which lies in the middle of color space, might be generated by flashes where the subject did not have enough information to make a confident guess about the color (red, green, blue). This could have been due to external noise in the stimulus (e.g. a stimulus mis-delivery, scattered light or random fluctuations in light intensity) or internal noise (i.e. neural). Additionally, our subjects may have adopted a strict criterion whereby they avoided calling a pinkish, less certain, flash ‘red’ and instead stingily labeled it ‘white’.

A preliminary experiment on a bright, yellowish background tested three locations in a single subject, who was given the choice of very red, weakly red, white, weakly green and very green. The weakly green category was never used and the subject did not feel confident in the usage of weak versus very red and indicated a preference for the simpler scheme used here^54^. In a second pilot experiment, the subject was permitted to respond arbitrarily along a continuous color wheel. However, the subject never felt intermediate colors were necessary to adequately describe the percepts on a white background and it was subsequently abandoned in favor of the five-choice paradigm.

### Background condition and colorimetry

Backgrounds were set with an external projector in Maxwellian view. The backgrounds consisted of the 840 nm imaging raster, the residual light from the 543 nm light source arising from an imperfect extinction of the optical switch used to control it, and the external projector illuminating the retina in Maxwellian view. In order to measure the chromaticity and luminance of the overall white and blue backgrounds engendered by these multiple sources, a patch the same size as and adjacent to the background was generated with the external projector. The subject was then given control over the chromaticity and luminance of the adjacent patch and asked to achieve a match in hue and brightness to the background. This procedure was carried out two times. The luminance and chromaticity of the match was then measured with a PR650 spectroradiometer (Photo Research). The chromaticity of the blue background was 0.16, 0.15 in CIE 1931 xy space and the luminance was 38 cd m^−2^. The white background had a chromaticity of 0.23, 0.30 and a luminance of 46 cd m^−2^. The photopic luminance of our backgrounds and the near foveal eccentricity probed reduced any contribution from rods to a minimum. The activity produced in L, M, S cones to each background was computed according to the transformation between CIE 1931 xy space and cone fundamentals^55^ (their Table 8.2.5). The quantal catches were then transformed into a MacLeod-Boynton color space constructed from the Stockman & Sharpe fundamentals^33^.

### Analysis

Responses from single cones were analyzed for percentage seen, color reported and dominant response. A cone’s dominant response corresponded to the color category that was elicited most frequently. Trials reported as ‘not seen’ were excluded from analysis, except where noted. Due to the similarity in trends between our two subjects, we analyzed the data from both subjects together, except where noted. All analyses were carried out in Matlab (Mathworks; Natick, MA). Matlab’s ‘knnsearch’ function was used to find the distance to the nearest S-cone and the nearest six neighbors. Regression and correlation analyses were conducted as described in the text using the ‘regstat’ and/or ‘corr’ functions in Matlab.

To compare whether L- or M-cones were more likely on average to generate a given color response, each cone was treated as a separate sample from the population of all L- and M-cones. For example, we sampled 159 L-cones and 99 M-cones on a blue background, found the percent of trials that each one of those cones elicited a ‘red’ response and then compared the distribution of percent red responses in our sample of L- versus M-cones. Therefore, the standard error was computed as the standard deviation divided by the square root of the number of cones and t-tests were used to asses the statistical significance between L- and M-cone distributions. One and two-sample t-tests were carried out on color naming data using the ‘ttest’ and ‘ttest2’ methods, respectively. Separately, we assessed the likelihood that a given color was equally distributed across L- and M-cone trials with Fisher’s Exact tests (Matlab’s ‘fishertest’). For a given color category, for instance blue, we first constructed a two-by-two contingency table. The total number of trials assigned to that category, blue in this example, were tallied for both L- and M-cone trials and recorded in each row of the first column. In the second column we tallied the total number of L- and M- trials assigned to categories other than blue. From the contingency table an odds ratio was computed, which indicated the likelihood of a blue report when an M-cone was targeted versus an L-cone. Finally, the probability that the observed distribution of responses occurred purely by chance was also computed.

The influence of the local neighborhood surrounding a targeted cone was analyzed by first finding the spectral type of the nearest six cones. We then computed the number (out of a possible 6) that were of opponent spectral sensitivity relative to the targeted cone. For example, an M-cone with 1 S-cone and 2 L-cones in its nearest six neighbors has 3/6 opponent neighbors. A Spearman rank correlation coefficient was used to analyze the statistical influence of opponent neighbors on color names, due to the ordinal nature of the opponent neighbor metric.

During each session, 6-10 cones were targeted 20 times each and the order of presentations were randomized to prevent subject bias. Subjects were permitted to take breaks as needed during the sessions. A subset of cones was targeted during at least two separate sessions (N=44) to test the repeatability of the percepts generated in the current paradigm. Two measures of repeatability were computed. First, we calculated for each cone the correlation coefficient (R) between the response distributions obtained during the two sessions. For example, a response vector of (6,14,0,0,0) (corresponding to # of white, red, green, blue, and yellow reports) and a second of (10,5,5,0,0) would produce a repeatability of 0.48 In the few cases when the same cone was targeted three or more times, R was computed between all combinations of days and the average was used as the repeatability of that cone. On the previously reported white background^23^ this analysis yielded a mean repeatability of 0.82 and a median of 0.91. On a blue background, the mean R value was 0.77 and the median was 0.94, which confirmed the reported percepts were highly repeatable. Secondly, we computed the percent agreement by finding the minimum number of times the subject reported each color, summing those and normalizing by N trials. For instance, in the example above, the subject agreed 6 + 5 + 0 + 0 + 0 = 11 times out of a possible 20, i.e. % agreement = 55.0.

It is worth noting that all of the above described analyses under-estimate the repeatability of color responses because mixing up a red and green sensation is more severe than switching a red and white percept. However, none of the analyses incorporated this feature of color vision. The fact that our subjects were very unlikely to switch between green and red suggests that the percepts were even more repeatable than captured by these analyses.

The datasets generated during the current study are available in the figshare repository, https://figshare.com/articles/AOSLO_single-cone_color_naming_data_files/3619713.

## Acknowledgments

This work was supported by an unrestricted grant from the Research to Prevent Blindness and NEI NIH grants R01EY023591, P30EY001730, P30EY003176, R21EY021642, R01EY021242, K23EY022412, T32EY07031 and F32EY027637. R.S. received support from the Fight for Sight Postdoctoral Award. W.S.T. received support from the American Optometric Foundation Ezell Fellowship. R.S. holds a Career Award at the Scientific Interfaces from Burroughs Wellcome Fund. J.N. is the Bishop Professor in Ophthalmology.

## Author contributions statement

All authors designed the research; B.P.S, R.S. and W.T. conducted the research; B.P.S. and R.S. analyzed the data and wrote the paper. All authors revised and approved the manuscript.

## Additional information

Competing interests A.R. has a patent (USPTO#7118216) assigned to the University of Houston and the University of Rochester which is currently licensed to Boston Micromachines Corp (Watertown, MA, USA). Both he and the company stand to gain financially from the publication of these results.

